# Protein aggregation capture assisted profiling of the thiol redox proteome

**DOI:** 10.1101/2024.12.21.629874

**Authors:** Ana Martinez-Val, Samuel Lozano-Juárez, Jorge Lumbreras, Irene Rodríguez, Marinela Couselo-Seijas, Ana Simón-Chica, Carlos Galán-Arriola, Rodrigo Fernández, Estefanía Núñez, Inmaculada Jorge, David Filgueiras-Rama, Borja Ibáñez, Jesús Vázquez

**Affiliations:** Cardiovascular Proteomics Laboratory, Centro Nacional de Investigaciones Cardiovasculares Carlos III (CNIC), Madrid, Spain; Centro de Investigación Biomédica en Red de Enfermedades Cardiovasculares (CIBERCV), Madrid, Spain; Centro Nacional de Investigaciones Cardiovasculares Carlos III (CNIC), Novel Arrhythmogenic Mechanisms Program, Madrid, Spain; Translational Laboratory for Cardiovascular Imaging and Therapy, Centro Nacional de Investigaciones Cardiovasculares Carlos III (CNIC), Madrid, Spain; Cardiovascular Health and Imaging Laboratory, Centro Nacional de Investigaciones Cardiovasculares (CNIC), Madrid, Spain; Instituto de Investigación Sanitaria del Hospital Clínico San Carlos (IdISSC), Cardiology Department, Madrid, Spain; Department of Cardiology, Hospital Universitario Clínico San Carlos, Madrid, Spain; IIS-Fundación Jiménez Díaz Hospital, Madrid, Spain

**Keywords:** Redox, proteomics, DIA, myocardial tissue, ischemia-reperfusion, atrial fibrillation

## Abstract

Oxidative damage is critical in various diseases, including cardiovascular and neurological conditions. Thiol redox reactions, acting as oxidative stress sensors, influence protein structure and function. Redox proteomics based on differential alkylation of reduced and oxidized Cys forms using mass spectrometry enables comprehensive analysis of thiol redox status in cells and tissues.

We introduce PACREDOX, an innovative redox proteomics approach based on the Protein Aggregation Capture (PAC) protocol and we demonstrate its compatibility with library free data-independent acquisition (DIA). PACREDOX reduces preparation time and costs compared to traditional methods, such as FASILOX, while maintaining thiol and proteome coverage. To enable library-free DIA, we corrected in silico spectral libraries in DIA-NN using experimental retention time data from beta-methylthiol-modified peptides. PACREDOX with DIA quantified 4,000 protein groups and ∼45,000 modified peptides in myocardial tissue from a porcine model of atrial fibrillation, including over 8,000 cysteine-containing peptides, 30% of which were reversibly oxidized.

Benchmarking PACREDOX and DIA against FASILOX in a myocardial infarction model reflects the potential and efficiency of this methodology to study oxidative damage. Overall, PACREDOX offers a high-throughput, cost-effective strategy for thiol redox proteome analysis, compatible with label-free quantitative workflows.

## Introduction

The study of redox post-translational modifications (PTMs) on cysteine thiols is essential to understand protein functionality and homeostasis. Redox PTMs, which alter proteins’ structural and functional properties, are produced when the thiol moiety in cysteine residues is modified as result of its participation in a redox reaction. Such reactions can either be induced by reactive oxygen species (ROS), or by specific enzymatic reactions. These modifications significantly impact cellular processes, as they regulate protein stability, activity, and interactions. Given the complexity of redox PTMs, redox proteomics has emerged as a valuable approach to map these modifications on a systems level. Mass spectrometry (MS) has become the primary tool for large-scale PTM analysis, allowing researchers to localize modifications at the amino acid level and to quantify relative abundance, thus providing comprehensive insights into redox-related protein modifications (1).

Redox PTMs arise when ROS interact with free thiols (SH groups) on cysteine residues, leading to a spectrum of oxidative modifications. Some of these modifications can be reverted by reductors and are so-called reversible. They include, among others, S-sulfenylation, S-nitrosylation, S-glutathionylation, disulfide bond formation, and S-polysulfidation. This reversibility enables dynamic regulation of protein function and contributes to redox signaling. On the other hand, irreversible modifications, like S-sulfinylation and S-sulfonylation, often result in permanent structural changes, marking proteins for degradation or altering their interactions with other molecules (2, 3).

In conventional bottom-up proteomics, proteins are usually reduced to optimize digestion with proteases and reversible Cys modifications are lost. Several experimental strategies have been developed to circumvent this problem to study redox PTMs. A popular strategy is differential alkylation, which involves selective reduction and tagging of different redox states. Initially, free thiol groups (corresponding to the basal or reduced Cys state) are blocked with an alkylating reagent to prevent further modifications. Subsequently, specific reductants are applied to the sample to reduce reversible oxidized forms, followed by a second alkylation step using a distinct reagent. This approach allows researchers to distinguish between different redox states and to tag specific modifications for further study.

For enhanced detection sensitivity, enrichment methods can be employed. OxICAT (isotope-coded affinity tags) employs thiol-reactive tags that facilitate both enrichment and quantification of cysteine modifications (4). Resin-assisted capture (RAC), which uses thiolated sepharose beads, covalently captures proteins containing free thiols. Another strategy, CysPAT, involves a thiol-reactive group with a linker and a phosphate group, which allows for protein trapping through immobilized metal affinity chromatography (IMAC) (5). Other techniques, such as GELSILOX and FASILOX, do not use enrichment steps but only require minor additions to conventional protocols and allow the analysis of different redox Cys states alongside overall protein alterations, making them versatile tools for redox proteomics (6, 7). In addition to these enrichment techniques, chemoselective probes are employed to detect specific redox PTMs with high specificity. These approaches involve the reaction of thiol redox forms with functionalized groups, such as biotin, which are then used to capture and enriched those modified peptides. Chemoselective probes have proven to be useful to study modifications such as S-nytrosylation (8) or S-persulfidation (9, 10). In contrast, direct detection approaches, which forego derivatization steps, have also emerged, allowing researchers to monitor redox PTMs with greater precision (11, 12).

In the recent years, aggregation of proteins into particles have been described to prepare samples for proteomics workflows, such as SP3 (Hughes, 2019, Nat Protocols) or PAC (13). These protocols not only increase the throughput of proteomics sample preparation, but in contrast to in-solution digestion strategies, they also clean up the sample. On the other hand, when compared to other sample preparation strategies such as filter-based, which also allow for solvent exchange, particle aggregation-based workflows are easier to streamline using magnetic racks and are not susceptible to delays in the protocol execution due to, for instance for FASP, filter blockage.

From a mass spectrometry perspective, redox proteomics has traditionally relied on data-dependent acquisition (DDA) for quantification. However, recent advances have seen data-independent acquisition (DIA) approaches gaining traction. Techniques like oxSWATH utilize DIA for comprehensive quantification in redox proteomics; however, this technique initially relies on DDA strategies to build spectral libraries required for DIA analysis (14). Although library-free methods remain largely unexplored in redox proteomics, they represent a promising area for future research. Computationally predicted libraries, such as DIA-NN (15) and Prosit (16), can predict IAM-modified peptides but are limited to this specific alkylating agent, restricting their application to broader redox PTM analysis. Emerging tools like DirectDIA in Spectronaut, which supports searches for various PTMs, offer potential solutions to this limitation, though they may not be accessible to all researchers due to licensing constraints.

In this work, we present an application of protein aggregation capture-based proteomics sample preparation to study the thiol redox proteome. Moreover, we present how it is possible to apply library free DIA analysis to boost the depth of the thiol redox proteome whilst reducing sample preparation timing and subsequent costs.

## Results

### Protein Aggregation Capture based differential alkylation for RedOx proteomics

In this work, we propose an adaptation of the Protein Aggregation Capture (PAC) digestion protocol (13) that supports differential alkylation of cysteines, named PACREDOX. To achieve this, we incorporated three additional steps into the standard PAC sample preparation workflow (**Figure 1A**). First, samples are homogenized, lysed, and proteins denatured in an SDS-containing buffer in the presence of iodoacetamide (IAM), which is used to carbamidomethylate free thiols. After boiling, samples are aggregated onto hydroxyl beads by adding acetonitrile. The buffer is then removed, and samples are reduced in a buffer containing dithiothreitol (DTT). Following reduction, the aggregation step is repeated with acetonitrile, and the buffer is removed again. The freshly created free thiols (corresponding to reversibly oxidized Cys) are then alkylated by incubating the beads with a buffer containing methyl-methanethiosulfonate (MMTS). After the second alkylation, proteins are aggregated once more onto the beads using acetonitrile. Importantly, skipping the aggregation after the reduction and second alkylation step resulted in significant loses of protein (**Suppl. Figure 1A**). Apart from these modifications, the PAC protocol proceeds without further alterations.

**Figure 1.**
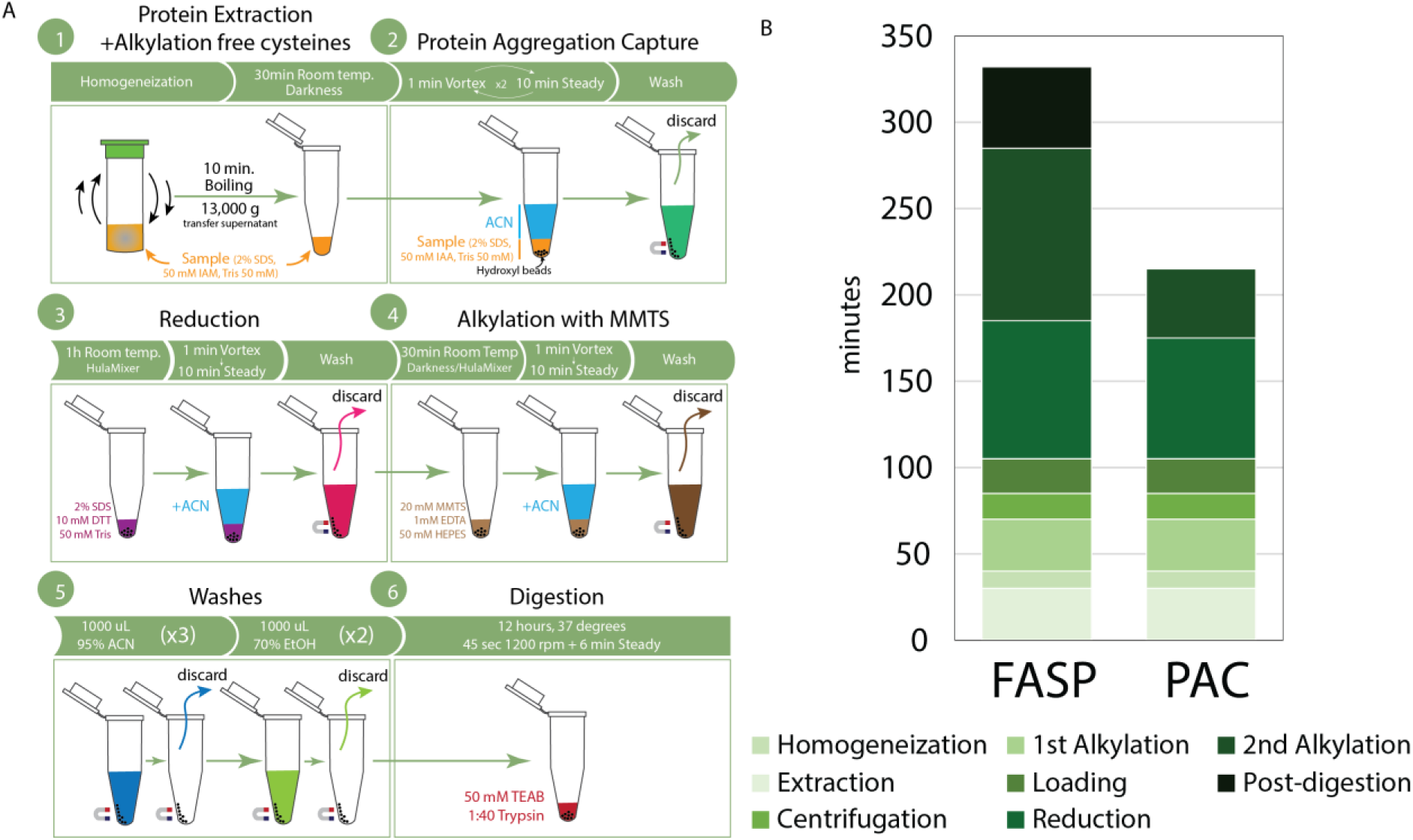
(A) Protein Aggregation Capture adapted workflow for differential alkylation of cysteine thiol groups to perform redox proteomics studies. (B) Stacked bar plot of expected times required for each step in FASP or PAC-based protocols for redox proteomics.

We compared this new protocol against the FASP-based differential alkylation protocol (FASILOX) previously developed by our group (6). The FASILOX protocol is based on the same differential alkylation strategy with IAM and MMTS, but it was originally designed to incorporate isobaric labelling for quantification. To assess the advantages of our PAC-based approach, we quantified the time required for differential alkylation and digestion when using PAC as described here versus FASP as outlined in the FASILOX protocol (**Figure 1B**). The centrifugation, washing and post-digestion steps in the FASILOX method significantly increase sample preparation time compared to the equivalent steps in PACREDOX. Additionally, it is important to note that this time estimate does not account for potential blockages of the FASP filters, which can further extend centrifugation times.

### Optimization of ‘in silico’ libraries for DIANN analysis of redox samples

To validate the usability of the PACREDOX protocol we analyzed whether the PAC-based differential alkylation is compatible with various quantification strategies. First, we first analyzed protein extracts from a porcine model for atrial fibrillation (AF) subjected to PACREDOX or FASILOX preparations and analysed using DDA, to validate that PACREDOX reported the same data as FASILOX. We observed that both PAC and FASP preparations resulted in equivalent depth of non-modified, IAM-labeled and MMTS-labeled peptides. The MMTS-peptides, corresponding to reversibly oxidized cysteines, made up about 30% of the reduced cysteine-containing peptides (**Supplementary Figure 1B**). That proportion aligns with the expected proportion in the tissues examined and with the accumulating evidence showing a potential role of oxidative stress and atrial cardiomyopathy in the pathogenesis of AF (17) and, particularly, the ratio of oxidized cysteine to reduced cysteine has been found increased in plasma samples of patients with AF (18).

Next, we evaluated whether differential alkylation with PACREDOX can be used to study the redox proteome through label-free quantification on data acquired using data-independent acquisition (DIA). Until now, redox proteomics has been largely limited to data-dependent acquisition (DDA) analysis due to the lack of in silico spectral libraries that encode peptides modified with secondary alkylation compounds, such as methyl-methanethiosulfonate (MMTS). Previous redox proteomics studies using DIA (9) have relied on experimentally-generated libraries. In this work, we sought to explore whether PACREDOX could be compatible with ‘in silico’ predicted libraries, such as those used in DIA-NN.

To assess this, we first evaluated the similarity between the fragmentation profiles of peptides labelled with IAM (i.e. carbamidomethylated peptides) and those labelled with MMTS (i.e. methylthiolated peptides). For this purpose, we analysed the PACREDOX DDA data obtained previously and evaluated whether the fragmentation profile of IAM-labelled and MMTS-labelled peptides was equivalent. To do so, we measured the correlation between the intensity of the fragments equivalent in IAM- and MMTS-labelled peptides. This analysis revealed that MMTS-labelling does not appreciably alter the fragmentation profile compared to IAM-labelled peptides (**Figure 2A-B**). In fact, the correlation of fragmentation profiles of the ∼400 peptide pairs analysed showed a median Pearson correlation value of 0.91, which increased to 0.94 when more than 12 fragments were available to calculate the correlation (**Figure 2B**). This result suggests that ‘in silico’ fragmentation predictions of IAM-labelled peptides can be translated to MMTS-labelled peptides.

**Figure 2.**
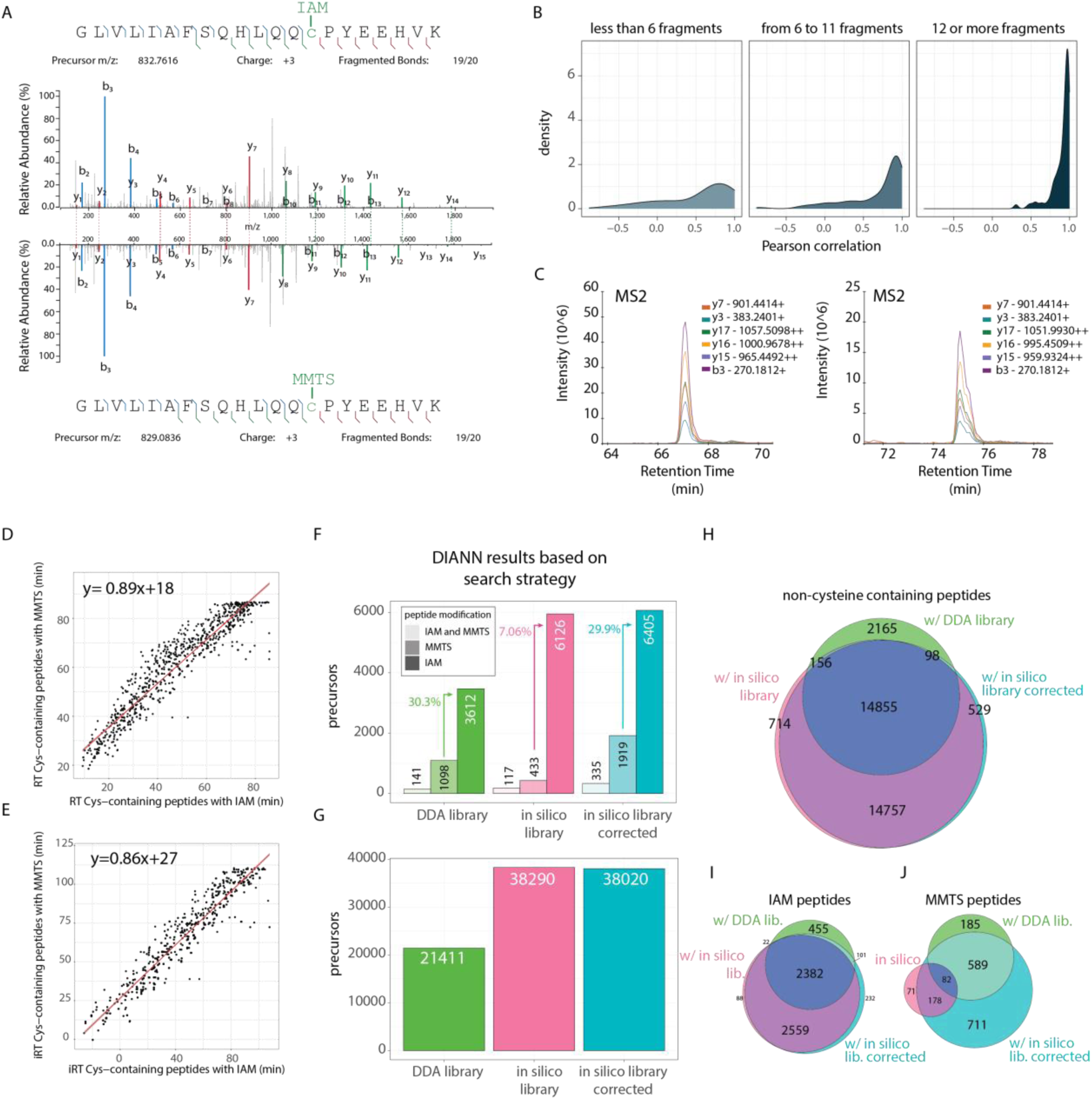
(A) Fragmentation spectrum of a cysteine-containing peptide labelled either with IAM (top) or with MMTS (mirrored, bottom). In blue: b-series fragments; in red: common y-series fragments and in green: specific y-series fragments. (B) Density plots of Pearson correlation values calculated between the fragmentation profiles (of equivalent fragment ion series) of IAM-labelled peptides and their MMTS-labelled counterparts, based on the number of fragments available for the comparison. (C) Extracted ion chromatograms at MS2 level for the two peptides from panel A (IAM-labelled on the left, MMTS-labelled on the right). (D) Scatter plot of representing retention times (RT) from IAM-labelled peptides vs their MMTS-labelled counterparts in an 88 minutes LC-run. Red line corresponds to the regression line, which equation is indicated in the top left corner. (E) Scatter plot representing iRTs from DIA-NN transformed library from IAM-labelled peptides vs their MMTS-labelled counterparts in the same analysis as in panel D. Red line corresponds to the regression line, which equation is indicated in the top left corner. (F-G) Bar plot of the number of cysteine-labelled precursors (F) or non-cysteine-containing precursors (G) identified in two samples in DIA-NN with three different search strategies: using a library generated in FragPipe from DDA files (green), using an ‘in silico’ library from DIA-NN (pink) and using an ‘in silico’ library from DIA-NN in which the retention times from MMTS-labelled peptides was corrected according to the equation from panel E (blue). (H-J) Euler diagrams showing the overlap between identified precursors in DIA-NN searches, either non-cysteine containing ones (H), IAM-labelled (I) or MMTS-labelled (J), when using either DDA-based library, DIA-NN ‘in silico’ library or DIA-NN ‘in silico’ library corrected for retention time.

However, when MMTS-labelled peptide spectra properties were calculated directly for the in-silico library used by DIA-NN, the rate of MMTS-labelled peptides was significantly lower than anticipated and the proportion of identified MMTS-labelled peptides with respect to IAM-labelled peptides was only 7% (**Figure 2F**). In contrast, when using an experimental DDA-based library, we observed that MMTS-labeled peptides, corresponding to reversibly oxidized cysteines, made up about 30% of the reduced cysteine-containing peptides (**Figure 2F**), which aligns with the expected proportion in the tissues examined and with the previous results obtained using DDA from samples generated with the FASILOX workflow (**Supplementary Figure 1B**).

Further examination of the results obtained using the DDA-experimental library revealed that MMTS-labelled peptides eluted consistently later their IAM-labelled counterparts (**Figure 2C-D**), presumably due to their higher hydrophobicity. We noticed that the iRTs of MMTS-labelled peptides could be inferred by a linear transformation of the iRT values of the IAM-labelled counterparts from DIA-NN libraries (**Figure 2D-E**). Therefore, we used this transformation to adjust the predicted retention times of MMTS-labelled peptides in the original ‘in silico’ library and reanalysed the samples using DIA-NN. Encouragingly, the iRT correction allowed us to recover the expected rate of reversibly oxidized cysteines to reduced cysteines (29.9%) (**Figure 2 F-G**), significantly enhancing the overall coverage of the thiol redox landscape in our DIA experiment without adversely affecting the identification of other peptides (**Figure 2H-J**). These findings indicate that it is possible to use prior knowledge for ‘in silico’ libraries to investigate modifications that were not initially implemented in the library prediction model. This adjustment demonstrates the viability of analysing the thiol redox proteome using DIA with ‘in silico’ predicted libraries.

Alternatively to DIA-NN, other strategies are available to study differential alkylation using DIA. On the one hand, some tools, such as Spectronaut, allow to generate pseudo-DDA data by deconvoluting DIA spectra, which can then be analysed using spectrum-centric search strategies. To test this, we analysed our data using Spectronaut (v19.2) with carbamidomethylation and beta-methylthiolation set as variable modifications on Cys. Spectronaut results reflected the expected proportion of 30% oxidized to reduced cysteines in our model (**Supplementary Figure 1A**). On the other hand, other algorithms based on library prediction, such as AlphaDIA (v1.8.2) are capable of training specific modifications (19), in our case for methylthiolated cysteines. Analysis of our samples using AlphaDIA trained for methylthiolated Cys resulted in slightly less identifications than Spectronaut or DIA-NN, but it still recapitulated the correct proportion of oxidized to reduced cysteines (**Supplementary Figure 1C**), indicating that AlphaDIA is also a good strategy for study differential alkylation in DIA datasets.

### PAC-redox using DIA recapitulates the oxidative patterns previously found in a pig model of ischemia-reperfusion injury

After demonstrating that PAC-based sample preparation can be combined with DIA to analyse the thiol redox proteome, we benchmarked this approach in a biological model of oxidative stress. We used myocardial tissue protein extracts from a pig model of ischemia-reperfusion (I/R) previously analysed using FASILOX (11) to assess the oxidative damage produced by reperfusion, and to evaluate the cardioprotective effect of preconditioning (20, 21). We compared samples from three pigs with no intervention in baseline conditions to those from three pigs subjected either to 40 minutes of ischemia followed by 90 minutes of reperfusion (120 min R), or from three pigs subjected to 120 minutes of ischemia without later reperfusion (120 min NOR). Additionally, we compared samples from three pigs subjected to ischemic preconditioning followed by 24 hours reperfusion (PRE 24h) and those from three non-preconditioned animals at 24 hours of reperfusion (**Figure 3A**).

**Figure 3.**
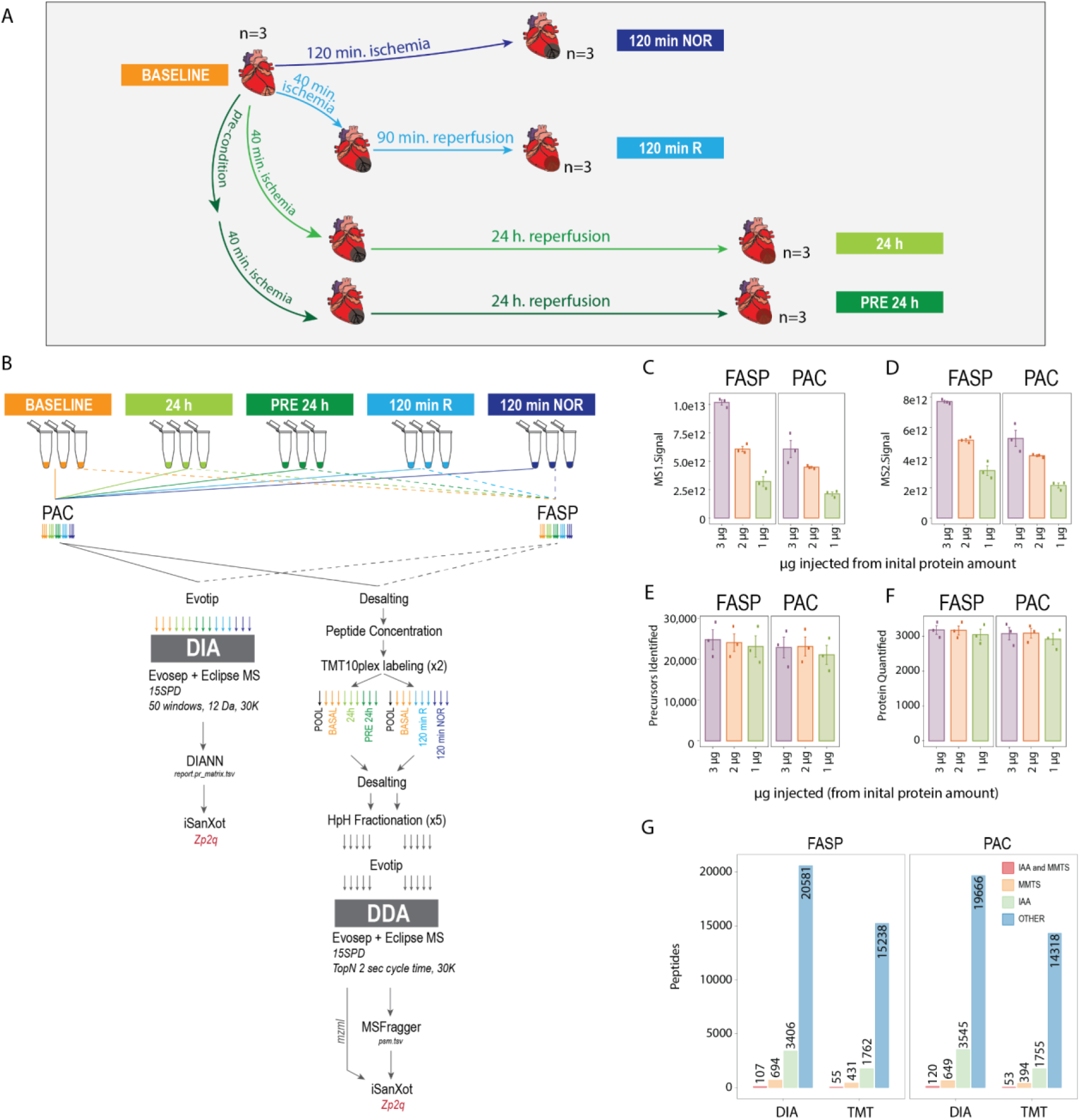
(A) Experimental design to study the effect of ischemia/reperfusion injury on cardiac tissue. (B) Sample preparation and data analysis workflow to study the redox proteome of heart extracts either using PAC or FASP-based digestion followed by either DIA label free or TMT quantification. (C-D) MS1 (C) and MS2 (D) signal measured in DIA analysis when injecting different expected amounts based on starting protein quantities in either FASP or PAC protocols. (E-F) Number of precursors (E) and proteins (F) identified using DIA analysis when injecting different expected amounts based on starting protein quantities in either FASP or PAC protocols. In C to F panels: height of the bar represents the average value and the error bar the standard error of the mean of 3 biological replicates. (G) Number of peptides used for quantification in PAC or FASP-based sample preparation and DIA label free or TMT workflows. Blue: non-cysteine containing peptides; green: cysteine-carbamidomethylated peptides; orange: cysteine-methylthiolated peptides and red: peptides containing both carbamidomethylated and methylthiolated peptides.

For a reliable benchmarking, we reanalysed the same protein extracts used by Binek et al., employing either PAC- or FASP-based digestion protocols, followed by quantification using either TMT-labelling or label-free DIA analysis (**Figure 3B**). The protein extracts were pre-processed as described in Binek et al., so free cysteines were already carbamidomethylated with IAM. Each sample began with 180 µg of protein, half of which (90 µg) was used in the PAC-based protocol and the other half in the FASP-based protocol. Both digestion protocols were conducted simultaneously in parallel.

From the sample preparation perspective, the label-free protocol was completed within two days (from digestion to MS injection), while the TMT-labelling protocol required two additional days for sample processing: after digestion, peptides underwent desalting and quantification before TMT labelling, followed by offline fractionation to obtain five fractions per sample. From the MS time perspective, label-free DIA sample data acquisition was completed in 24 hours using a 15 SPD gradient on an Evosep One. On the other hand, each TMT-labeled experiment, consisting on two TMT 10 plex of 5 fractions and 1 non-fractionated sample each, took a total of 19.2 hours using the 15 SPD gradient as well. Overall, DIA based MS aquisition required 1.6 hours per sample, whilst one TMT10plex-labeled sample (9 samples and 1 internal standard) resulted in 1.06 hours per sample, although since our design required to use two TMT10plex, the actual time analysis per sample increased to 1.28 hours. The time required for MS analysis either using DIA or TMT will vary depending on the experimental design. For instance, TMT-based protocol could have reduced the MS acquisition time by using TMTpro agents to include all replicates in a single experiment, requiring only one fractionation (see Discussion).

The first comparison between PAC and FASP digestion methods focused on recovery efficiency (i.e., the proportion of peptides recovered given a starting protein amount). Peptides were collected immediately after quenching digestion, and different proportions were loaded into Evotips assuming 100% recovery efficiency from protein starting amount. These samples were analyzed with DIA, and raw files were processed using DIA-NN independently for each condition. We observed that FASP yielded higher MS1 and MS2 signal intensities than PAC (**Figure 3C-D**). Specifically, injection of an estimated amount of 2 µg from FASP-prepared samples yielded comparable intensity to 3 µg from PAC samples, indicating that the FASP protocol incurs fewer peptide losses (**Figure 3C-D**). Despite this, protein and precursor identification performance did not significantly differ between PAC and FASP samples (**Figure 3E-F**). We further compared the identification rate of all combinations of sample preparation and MS data acquisition methods. We found that label-free DIA had consistently higher identification rate than TMT for a similar MS acquisition time (**Figure 3G**). However, identification performance using TMT could be enhanced by using a more comprehensive offline fractionation scheme or longer gradients, although that might result in higher cost and complexity. Apart from this the performance was similar independently on the digestion method of choice for both quantification strategies (**Figure 3G**). Overall, these results support the validity of PACREDOX combined with DIA as an alternative to more elaborate protocols such as FASILOX.

We next assessed quantification performance for PAC and FASP-based protocols coupled to either label-free DIA analysis and the original FASILOX workflow (i.e. FASP coupled to isobaric labeling). The quantitative data at the scan (for TMT) or precursor (for DIA) levels were processed by the iSanXoT package (22, 23), using the standard workflow for the analysis of post-translational modifications (16). The quantitative values were expressed as log_2_-ratios, using as reference for the denominator the average of the baseline condition samples. The quantitative scan (for TMT) or precursor (for DIA) values were then integrated to peptide values (pr2p), and the peptide values to protein values (p2q), to obtain standardized peptide values *Zp2q*, which are independent from protein changes (16). iSanXoT uses the Generic Integration Algorithm (GIA) (24) to model the distribution of variances in each integration. We observed that while the global variance at peptide (p2q) and protein (q2all) was clearly higher in DIA data than that of TMT (**Supplementary Figure 2A-B**), irrespective of the sample preparation protocol used (**Supplementary Figure 2A**). This agrees with the lower accuracy of label-free quantification in comparison with stable isotope labelling quantification, where the intensities of the different samples are measured within the same spectrum.

In order to study the thiol redox proteome, we analysed the distribution of peptide values for three separate populations: reversibly oxidized Cys-containing peptides, reduced Cys-containing peptides, and non-Cys-containing peptides. The *Zp2q* values from the peptides without Cys and the reduced Cys-peptides followed very accurately the theoretical N(0,1) distribution (**Figure 4A-C**) both for FASILOX and for the PACREDOX-DIA approach, demonstrating that the results obtained with the PACREDOX protocol, like those with FASILOX (6), can be accurately modelled using the statistical GIA framework. In contrast, the distribution of oxidized Cys-containing peptides deviated from the null hypothesis, showing increased levels in the samples subjected to ischemia-reperfusion for 120 minutes and 24 hours (**Figure 4A-C**) both in FASILOX and PAC-DIA data, reflecting oxidative thiol damage, as previously described (7). Notably, in the absence of reperfusion the oxidative damage after 120 minutes was clearly lower than that observed with 90 minutes reperfusion using both preparation protocols (**Figure 4A-B and E** and **Supplementary Figure 2C**). Similarly, the data obtained by the two approaches also showed that ischemia preconditioning mitigated the oxidation observed after 24 hours of I/R (**Figure 4C-E** and **Supplementary Figure 2C**). All these results indicate that neither the choice of digestion strategy nor the quantification strategy alter the outcomes of the experiment, which reflect the same biological effects.

**Figure 4.**
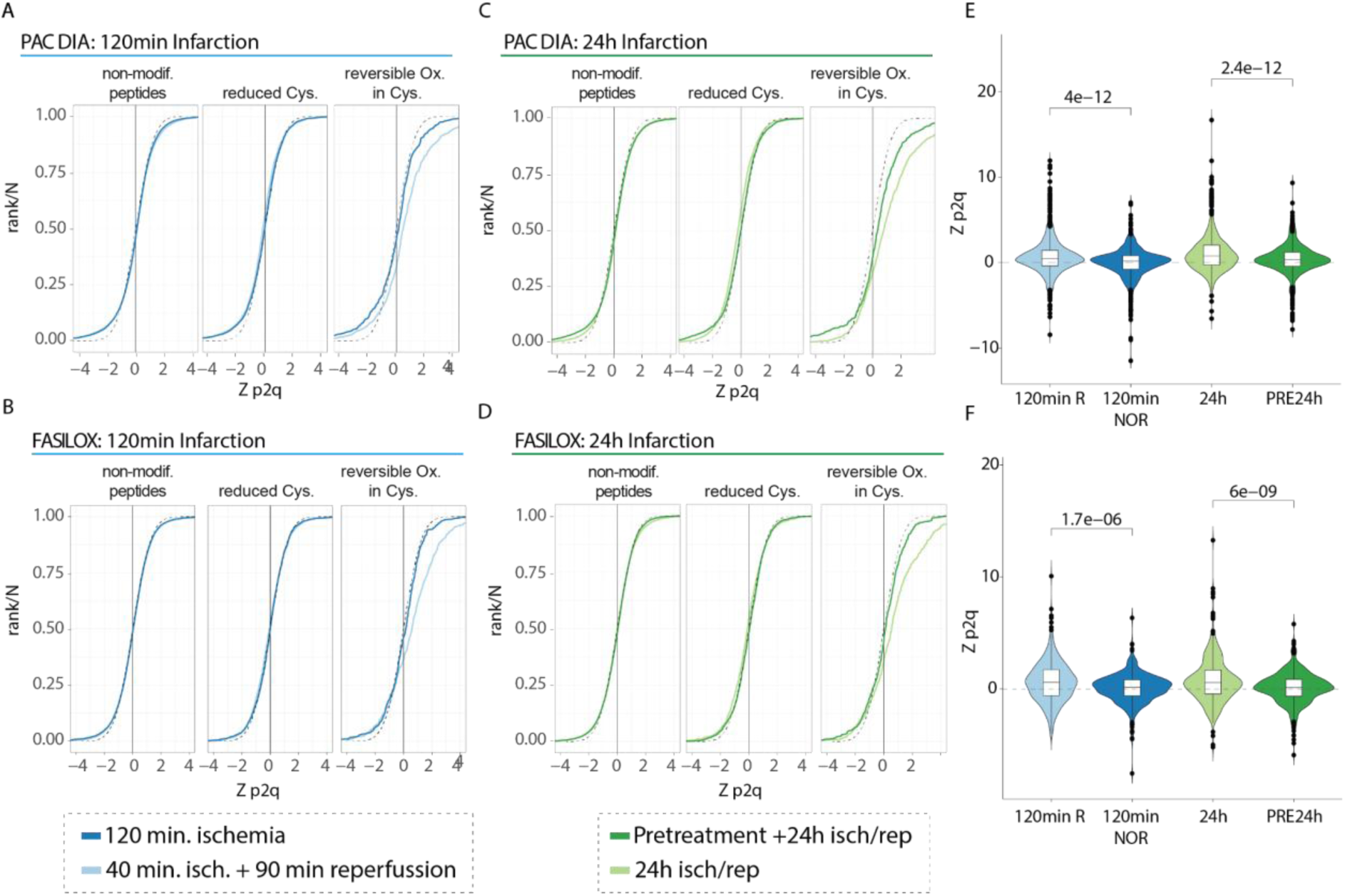
(A-D) Sigmoid curves showing the ranked distribution of Z values from the integration peptide to protein (p2q) for three peptide populations: non-cysteine containing peptides, reduced cysteines and reversible oxidized cysteines. (A) Sigmoid curves in PACREDOX-DIA redox experiment for 120 minutes of I/R. (B) Sigmoid curves in FASILOX experiment for 120 minutes of I/R. In A and B, light blue corresponds to 40 minutes ischemia + 90 minutes of reperfusion (120min R) and dark blue corresponds to 120 min of ischemia without reperfusion (120min NOR). (C) Sigmoid curves in PACREDOX-DIA redox experiment for 24 hours of I/R. (D) Sigmoid curves in FASILOX experiment for 24 hours of I/R. In C and D, light green corresponds to 40 minutes ischemia + 24 hours of reperfusion (24h) and dark green corresponds to 40 minutes ischemia + 24 hours of reperfusion on preconditioned pigs (PRE24h). In all panels, the black dotted line represents the normal distribution N(0,1). (E) Boxplot and violin plot of the Zp2q values for MMTS-labeled peptides in the PAC-DIA redox experiment for the different conditions, all relative to baseline samples. On top, p-value from a two-sample t-test. (F) Boxplot and violin plot of the Zp2q values for MMTS-labeled peptides in the FASILOX experiment for the different conditions, all relative to baseline samples. On top, p-value from a two-sample t-test.

However, it is relevant to indicate that DIA achieved higher coverage than TMT of the thiol redox proteome. Whilst DIA provided evidence of oxidized disulfide bonds in 177 protein-coding genes (**Supplementary Figure 3A**), TMT covered only 90 protein-coding genes with reversibly-oxidized cysteines (**Supplementary Figure 4A**). By functionally annotating those proteins we can gain a further understanding of the pathways that might be affected by redox reactions. There were four main functions that grouped most of these proteins: cellular respiration, humoral immune response, cell adhesion and sarcomere organization. However, the coverage of those pathways was more extensive in DIA derived data (**Supplementary Figures 3B and 4B**). Moreover, thanks to DIA we can identify proteins that can be relevant in the I/R injury process that were not identified by TMT. Such is the case of Myomesin 2 (MYOM2), which is a major component of the vertebrate myofibrillar M band which has been described to be oxidized in heart failure (25). Only in DIA we have evidence of four reversibly oxidized cysteine-containing peptides, which show the expected effect of increased oxidation after I/R injury (**Supplementary Figure 2D**).

Finally, we assessed technical reproducibility in DIA between PAC or FASP-based workflows. We already showed that both strategies report a similar number of peptides, but FASP has better recovery. From a quantification perspective, however, we did not observe relevant differences between both strategies when using DIA (**Supplementary Figure 5A-B**) and the quantification of reversibly-oxidized peptides correlated well between both digestion strategies (**Supplementary Figure 5C**).

## Discussion

In this study, we introduced a novel sample preparation strategy leveraging the Particle Aggregation Capture approach for protein digestion, enabling differential alkylation and facilitating in-depth analysis of the thiol redox proteome. This methodology significantly reduces sample processing time, which is crucial for increasing experimental throughput and potentially minimizing variability associated with complex preparation techniques.

Additionally, we demonstrated that this kind of thiol redox proteomics, originally proposed for isobaric-labelling, can be performed using label-free strategies and is compatible with data-independent acquisition (DIA) without requiring experimental libraries. Notably, we utilized predicted libraries that can be adjusted to accommodate additional modifications beyond their initial training.

Our findings confirm that DIA can achieve a deeper proteome coverage compared to data-dependent acquisition (DDA) methods, such as tandem mass tag (TMT) labeling, given similar acquisition times. While a comparable depth to DIA might be achievable in TMT by increasing the number of fractions analyzed, this would substantially raise sample processing times. From a quantification standpoint, DIA data displayed greater variability than TMT, an important consideration in thiol redox proteomics where quantification is peptide-based. This variability must be considered when designing experiments; achieving comparable statistical power with DIA as with TMT may require additional replicates to balance the inherent quantification variability in DIA. Finally, our work shows for the first time that the Generic Integration Algorithm, applicable through the iSanXoT package, is a valid model for the accurate analysis of DIA data.

In summary, redox proteomics, driven by advances in mass spectrometry and computational tools, offers powerful methodologies for understanding redox PTMs on cysteine residues. Through a combination of differential alkylation, enrichment strategies, and evolving MS techniques, redox proteomics continues to describe the complex redox landscape in cellular systems. While challenges remain, particularly regarding library-free approaches and tool accessibility, ongoing innovations such as the one presented in this work promise to expand our understanding of protein modifications and their impact on cellular function.

## Ethics statement

All animal procedures related to the atrial fibrillation porcine model were approved by the Comunidad de Madrid (Ref# PROEX097/17 & PROEX078.8/21) and conformed to the regulations outlined in the EU Directive 2010/63EU and Recommendation 2007/526/EC regarding the protection of animals used for experimental and other scientific purposes.

The ischemia/reperfusion study was approved by the Ethics Committee of the Conserjería de Medioambiente, Ordenación del Territorio y Sostenibilidad de la Comunidad de Madrid (PROEX51/13, 3 February 2013; PROEX034/19, 23 October 2019; and PROEX 176.3/20, 23 July 2020).

## Authors contribution

A.M.-V. performed all MS experiments, analysed the data and prepared the manuscript. S.L. performed data analysis. I.R. and J.L. performed the proteomics sample preparation. M.C-S, A.S-C and D.F-R. generated the atrial fibrillation porcine model. C.G., R.F. and B.I. generated the ischemia/reperfusion injury porcine model. A.M-V., E.N., I.J. and J.V. devised the project, planned the experiments and revised the manuscript. A.M-V., D.F-R., B.I. and J.V. provided funding. All authors contributed to and approved the final version of the manuscript

## Acknowledgements

We thank all members of the Proteomics Unit and Cardiovascular Proteomics Groups from CNIC for discussions regarding this work. A.M-V. work is funded by the grant “Ayudas de Atracción de Talento Investigador César Nombela 2023” (2023-T1/SAL-GL-28990, Comunidad de Madrid). J.L. work is funded by the grant “Ayudas para la contratación de ayudantes de investigación y técnicos de laboratorio 2022” (PEJ-2021-TL/BMD-23107, Comunidad de Madrid). I.R. work is funded by the grant “Ayudas para la contratación de ayudantes de investigación y técnicos de laboratorio 2023” (PEJ-2023-TL/SAL-GL-26599, Comunidad de Madrid) and the European Social Fund Plus (2021-2027). This study was supported by competitive grants PID2021-122348NB-I00 funded by MICIU/AEI/ 10.13039/501100011033 and by “ERDF A way of making Europe”, PLEC2022-009298, PLEC2022-009235 and EQC2021-007053-P funded by MICIU/AEI/10.13039/501100011033 and by “European Union NextGenerationEU/ PRTR”, and S2022/BMD-7333-CM (INMUNOVAR-CM) funded by Comunidad de Madrid. The project leading to these results has received funding from “la Caixa” Foundation under the project code LCF/PR/HR22/52420019. The CNIC is supported by the Instituto de Salud Carlos III (ISCIII), the Ministerio de Ciencia, Innovación Y Universidades (MICIU) and the Pro CNIC Foundation), and is a Severo Ochoa Center of Excellence (grant CEX2020-001041-S funded by MICIU/AEI/10.13039/501100011033). Research in DFR laboratory was supported by the European Union Horizon 2020 research and innovation program under Grant Agreement#965286, and by Program S2022/BMD-7229 -CM ARCADIA-CM funded by Comunidad de Madrid.

## Materials and Methods

### Reagents and Tools table

| Reagent/resource | Reference or source | Identifier or catalog number |
| --- | --- | --- |
| SDS 20% w/w | Bio-Rad | 1610418 |
| Tris | Cytiva | 17132101 |
| Clorhidric acid (HCl) | Merck | 1.003171000 |
| Water (H<sub>2</sub>O) | Fisher Scientific | 10728098 |
| Dithiothreitol (DTT) | Plus One | 17-1318-02 |
| Iodoacetamide (IAM) | Merck | I6125 |
| Methyl methanethiosulfonate (MMTS) | Fisher Scientific | 23011 |
| MagResyn Hydroxyl Beads | Resyn Biosciences | MR-HYX050 |
| Acetonitrile | Merck | 1.000292500 |
| Ethanol | Merck | 1.117272500 |
| Ammonium bicarbonate | Sigma-Aldrich | A6141 |
| Triethylammonium bicarbonate | Fisher Scientific | 90114 |
| Sequencing Grade Modified Trypsin | Promega | V5111 |
| Trifluoroacetic acid | Fisher Scientific | 85183 |
| RC DC™ Protein Assay Kit II | BioRad | 5000122 |
| Urea | Cytiva | 17131901 |
| Sodium chloride | Merck | 1.06404 |
| Thermo Scientific™ Conjunto de reactivos de etiqueta isobárica TMT10plex™ | Fisher Scientific | 15285743 |
| Kit de fraccionamiento de péptidos de fase inversa de pH alto Pierce™ | Fisher Scientific | 84868 |
| Nanosep Centrifugal Devices with Omega Membrane | Pall Life Sciences | OD303C34 |
| Oasis HLB 1 cc Vac Cartridge, 10 mg Sorbent per Cartridge, 30 µm, 100/pk | Waters | 186000383 |
| Evotip Pure 10x96 tips | Evosep | EV2013 |
| Formic acid | Merck | 5.33002 |
| 2-Propanol | Honeywell | 34965 |
| HEPES | Sigma-Aldrich | H4034 |
| Ethylenediaminetetraacetic acid (EDTA) | Plus One | 17-1324-01 |
| Ammonia solution 25% | Merck | 1.05432 |
| Endurance column 15 cm 150x19 | Evosep | EV1106 |
| ID 30 µm stainless steel emitter | Evosep | EN1086 |

### Methods and Protocols

#### Experimental design and animal models

##### Porcine model of long-lasting lone persistent atrial fibrillation (AF)

Yucatan-Large White crossbred pigs were used to generate the long-lasting lone AF model as previously described (17). Left and right atrial samples were taken at the time of euthanasia from left atrial free wall and from the right atrial appendage.

##### Porcine model of ischemia/reperfusion (I/R)

Castrated male Large White pigs weighing 30–40 kg were used for the pig I/R experiments as described (11). Briefly, closed-chest, 40 min mid-left anterior descending (LAD) coronary artery occlusion was performed and animals were sacrificed after 90 minutes and 24 hours of reperfusion. Additional animals were sacrificed 120 minutes after artery occlusion without reperfusion and another group of animals were subjected to ischemic preconditioning prior to I/R and sacrificed after 24 hours of reperfusion. Other pigs were sacrificed with no intervention other than baseline cardiac magnetic resonance (CMR) and served as controls. Animals were immediately euthanized after the last follow-up CMR scan coinciding with indicated time-points, and transmural myocardial tissue samples from the ischemic area were rapidly collected for proteomics, frozen in liquid nitrogen and stored at -80 degrees until proteomics processing.

#### Tissue sample preparation for Proteomics Analysis of differential cysteines alkylation following FASP digestion method

Protein extracts from homogenized tissue were obtained using ceramic beads (MagNa Lyser Green Beads apparatus, Roche, Germany) in extraction buffer (50 mM Tris-HCl, 1 mM EDTA, 1.5% SDS, 50 mM iodoacetamide (IAM), pH 8.5). Protein concentration was quantified using RC DC Protein Assay (Bio-Rad). Samples were subjected to tryptic digestion using the filter-assisted sample preparation protocol (FASP) according to the previously published method. Briefly, 90 µg of protein extract was diluted in urea sample solution (8 M urea in 100mM Tris– HCl, pH 8.5) and loaded on filters (30KDa pore size, Nanosep) by centrifugation (10 minutes at 11,000 g). Samples were washed with 7.5 M urea in 100mM Tris–HCl (pH 8.5). Protein thiol groups were reduced with 10 mM dithiothreitol (DTT) in 7.5 M urea in 100mM Tris–HCl (pH 8.5) for 30 minutes at 56 degrees with gentle agitation. Samples were washed with 50 mM HEPES 1 mM EDTA (pH 7.5), and then alkylated using 50 mM methyl methanethiosulfonate (MMTS), 50 mM HEPES, 1 mM EDTA (pH 7.5) for 30 minutes at room temperature with gentle agitation and protected from light. Samples were washed three times with 7.5 M urea in 100mM Tris–HCl, pH 8.5 and then three times with 100 mM ammonium bicarbonate (pH 8.5). Protein digestion was carried out overnight at 37°C with sequencing grade trypsin (Promega, Madison, WI, USA) at 1:40 (w/w) enzyme:protein ratio in digestion buffer (50 mM ammonium bicarbonate, pH 8.5), after which the resulting tryptic peptides were recovered by centrifugation. Trifluoroacetic acid was added to a final concentration of 1% and the peptides were desalted on C18 Oasis HLB extraction cartridges (Waters Corporation, Milford, MA, USA) and dried-down.

#### Tissue sample preparation for Proteomics Analysis of differential cysteines alkylation following PAC digestion method

Protein extracts from homogenized tissue were obtained using ceramic beads (MagNa Lyser Green Beads apparatus, Roche, Germany) in extraction buffer (50 mM Tris-HCl, 1 mM EDTA, 1.5% SDS, 50 mM iodoacetamide (IAM), pH 8.5). Protein concentration was quantified using RC DC Protein Assay (Bio-Rad). Briefly, to the volume equivalent 90 µg of protein, 100% of acetonitrile was added to achieve a final concentration of 70% of acetonitrile, followed by 10 µL of hydroxyl beads (Resyn Biosciences). Samples were aggregated by two steps of vortexing for 1 minute followed by waiting for 10 minutes. Samples were placed on a magnetic rack, and supernatant was discarded. Samples were incubated for 30 minutes at 56 degrees with 100 µL of reducing buffer (50 mM Tris (pH 8.5), 10 mM DTT, 2% SDS) mixing at 350 rpm. 233 µL of 100% of acetonitrile was added to the sample, followed by vortexing for 1 minute and resting for 10 minutes. Samples were placed back on a magnetic rack, and supernatant was discarded. Samples were incubated for 30 minutes at room temperature with 100 µL of secondary alkylation buffer (20 mM MMTS, 50 mM HEPES 1 mM EDTA, pH 7.5) mixing at 350 rpm. 233 µL of 100% of acetonitrile was added to the sample, followed by vortexing for 1 minute and resting for 10 minutes. Samples were placed back on a magnetic rack, and supernatant was discarded. Without removing the samples from the magnetic rack, samples were washed three times with 1 mL of 95% acetonitrile and two times with 1 mL of 70% ethanol. After removing the last wash, the digestion was performed overnight in 50 µL of 50 mM of TEAB containing sequencing grade trypsin (Promega, Madison, WI, USA) in ratio 1:40 (w/w, enzyme:protein). Trifluoroacetic acid was added to a final concentration of 1% and the peptides were desalted on C18 Oasis HLB extraction cartridges (Waters Corporation, Milford, MA, USA) and dried-down.

#### Labelling of peptides using TMT reagents followed by off-line HpH fractionation

Peptides were resuspended in 40 µL of 100 mM TEAB and peptide concentration was quantified by Direct Detect IR spectrometer (Millipore, Billerica, MA, USA). Peptides were subjected to multiplexed isobaric labelling using two tandem mass tag 10plex (TMT10plex) kits following manufacturer instructions. Each TMT 10plex batch was used to label 9 samples and one channel was reserved for reference internal standard samples created by pooling the four samples. Labelling scheme for TMT #1 was as follows: 126: Pool, 127N to 128N: Baseline, 128C to 129C: 120 minutes ischemia/reperfusion, 130N to 131N: 120 minutes ischemia. Labelling scheme for TMT #2 was as follows: 126: Pool, 127N to 128N: Baseline, 128C to 129C: 24 hours of ischemia/reperfusion, 130N to 131N: preconditioning followed by 24 hours of ischemia/reperfusion. Baseline samples used in TMT #1 and TMT #2 can be considered equivalent, since both proceed from the sample digestion and labelling step. TMT labelled peptides were mixed, dried down to remove the acetonitrile and desalted on C18 Oasis HLB extraction cartridges (Waters Corporation, Milford, MA, USA). Desalted peptides were dried down and resuspended in 400 µL of 0.1% TFA, 40 µL was used for single-shot analysis of the unfractionated sample and the rest was fractionated into five fractions using the high pH reversed-phase peptide fractionation kit (Thermo Scientific).

#### LC-MS/MS analysis

Samples were analyzed using an Endurance Evosep column (15 cm–150 μm-C18 1.6 μm) interfaced with the Orbitrap Eclipse Tribrid Mass Spectrometer (Thermo Scientific) using an EasySpray Ion Source with an in-house fabricated column oven to maintain the temperature at 55 °C. In all samples, spray voltage was set to 2.1 kV, funnel RF level at 40, and heated capillary temperature at 310°C. Samples were separated on an Evosep One LC system using the pre-programmed gradient for 15 samples per day (SPD).

For analysis using DIA, full MS resolutions were set to 120,000 at m/z 200 and the full MS AGC target was 300% with a maximum injection time (IT) of 45 ms. The AGC target value for fragment spectra was set to 1000%. 50 windows of 12 Th scanning from 400 to 100 m/z were employed with an overlap of 1 Da. MS2 resolution was set to 30,000, IT to 54 ms, and normalized collision energy (NCE) to 30 %.

For analysis of TMT samples using DDA, the MS was operated at a full MS resolution of 120,000 with a full scan range of 375 to 1500 *m/z*. The full scan AGC target was set to 100%. During a fixed cycle time of 2 second, precursors were isolated with a window of 1.2 Th and fragmented using HCD with 36 % NCE. Fragment ion scans were recorded at a fixed resolution of 30,000 and with a maximum IT of 54 ms and an AGC target value of 100%.

For analysis of comprehensive DDA samples, LC-MS analysis was done using an Easy nanoLC 1000 (ThermoFisher Scientific) coupled to an Orbitrap Fusion Tribid Mass Spectrometer (Thermo Fisher Scientific) using an Acclaim PepMap 100 C18 2 cm x 75 μm ID as trapping column (Thermo Fisher Scientific) and a PepMap RSLC C18 EASY-Spray column 50 cm x 75 μm ID as analytical column (Thermo Fisher Scientific). Peptides were loaded in buffer A (0.1% of formic acid in water (v/v)) and eluted with a 270 min linear gradient of buffer B (100% ACN, 0.1% formic acid (v/v)) at 200 nl/min. Mass spectra were acquired in data-dependent manner, with an automatic switch between MS and MS/MS with a “Top-15” method and 40 s dynamic exclusion. MS spectra were acquired in the Orbitrap analyser using full ion-scan mode with a 390-1700 m/z range and 60,000 FT resolution. The automatic gain control target was set at 1 x 106 with 50 ms maximum injection time. HCD fragmentation was performed at 30% of normalized collision energy and MS/MS spectra were analysed at a 30,000 resolution in the Orbitrap with automatic gain control and 100 ms maximum injection time.

#### Peptide and Protein Identification of TMT datasets

TMT raw files were analyzed using MSFragger (v4.1) in FragPipe (v22) using the Sus Scrofa proteome (UP000008227, accessed 25-03-2024). Fixed modification of the TMT label was set of lysine and peptide N-terminal. Variable modifications used were: TMT label on serine, oxidation on methionine, carbamidomethylation and beta-methylthiolation on cysteine and acetylation on protein N-terminal. Enzyme/cleavage rule was set to trypsin and a maximum of 3 variable modifications per peptide were allowed. Post-analysis validation and FDR control was performed using Percolator, whilst protein inference was performed using ProteinProphet with default parameters in FragPipe. TMT reporter intensities were extracted from mzML transformed raw files and corrected for isotopic impurities in iSanXoT (v1.2.13).

#### DDA-based library generation

DDA raw files were analyzed using MSFragger (v4.1) in FragPipe (v22) using the Sus Scrofa proteome (UP000008227, accessed 25-03-2024). Variable modifications used were: oxidation on methionine, carbamidomethylation and beta-methylthiolation on cysteine and acetylation on protein N-terminal. Enzyme/cleavage rule was set to trypsin and a maximum of 3 variable modifications per peptide were allowed. MSBooster was enabled. Post-analysis validation and FDR control was performed using Percolator, whilst protein inference was performed using ProteinProphet with default parameters in FragPipe. Spectral library generation was enabled with default values. Resulting library file was employed for DDA-based library searches in DIA-NN.

#### Peptide and Protein Identification of DIA datasets

DIA raw files were analyzed using DIA-NN (v 1.8.1) using an ‘in silico’ predicted library from the Sus Scrofa proteome (UP000008227, accessed 25-03-2024). For library prediction the enzyme/cleavage rule was set to trypsin, one missed cleavage per peptide and two variable modifications per peptide were allowed. Variable modifications used were IAM and MMTS on cysteine and oxidation on methionine. MMTS retention times were corrected based on experimental data as follows: iRTs from the DIA-NN spectral library for MMTS containing peptides were corrected using the following equation:

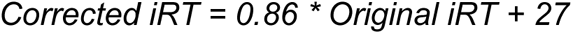

Cross-run normalization was turned-off and remaining options were set as default.

#### Peptide and protein quantification

The quantitative information derived from TMT reporter intensities and DIA precursor intensities was integrated from the spectrum level (for TMT data) or precursor (for DIA data) to the peptide level (“pr2p”) and subsequently to the protein level (“p2q”), according to the WSPP model (Navarro et al 2013) and the Generic Integration Algorithm (GIA) (Garcia-Marques et al 2016), using the iSanXoT program version 1.2.13 (Trevisan-Herraz et al 2019, Rdoriguez et al 2024). In this model, quantitative protein values are expressed using the standardized variable *Z* (i.e., normalized log2-ratios expressed in units of standard deviation according to the estimated variances).

## Supplementary Data

**Supplementary Figure 1.**
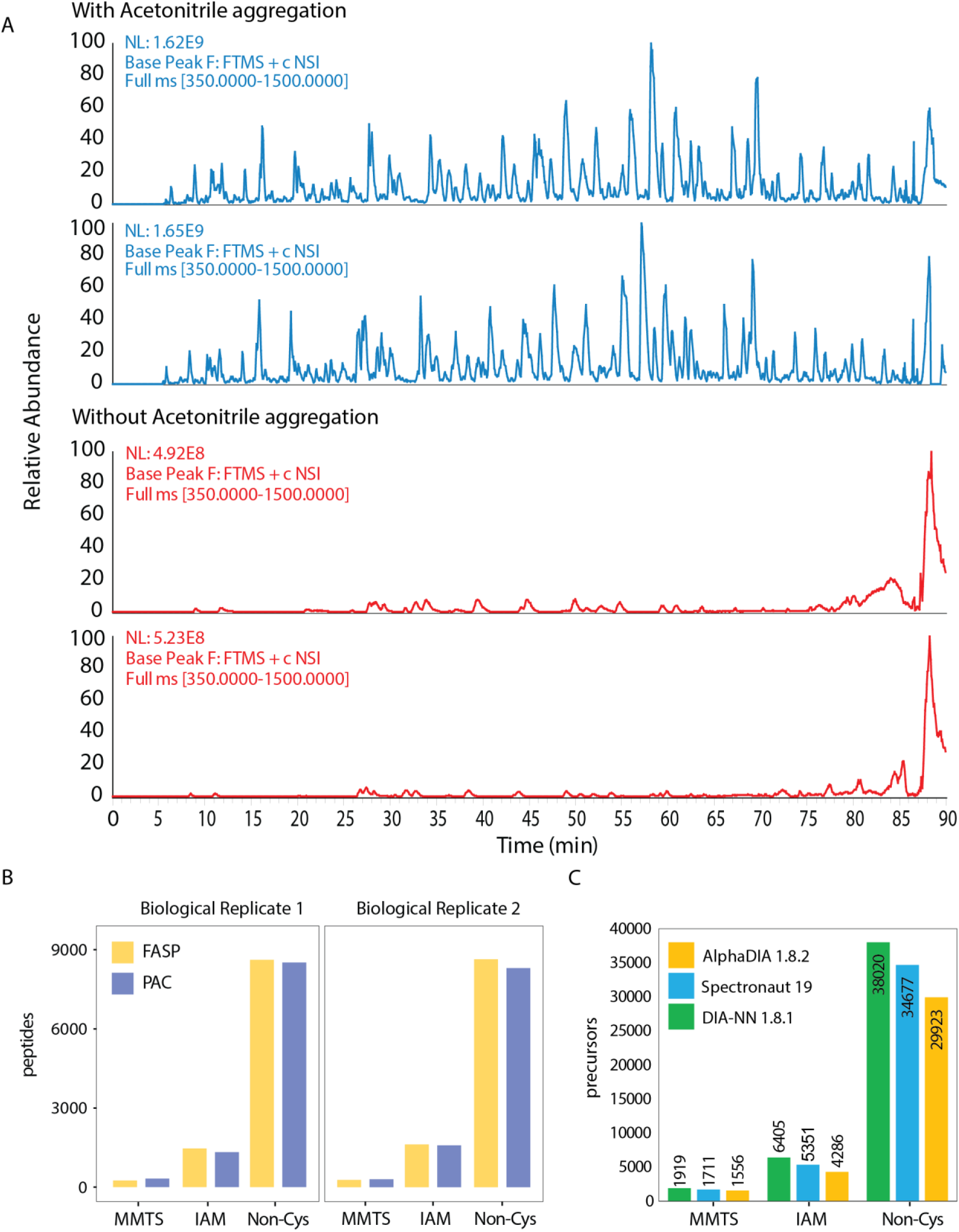
(C) Number of identified precursors in two raw files from porcine myocardial tissue processed using the PAC-redox protocol in three different DIA search engines without the use of experimental libraries: green - DIA-NN 1.8.1, blue - Spectronaut 19, yellow - AlphaDIA 1.8.2.

**Supplementary Figure 2.**
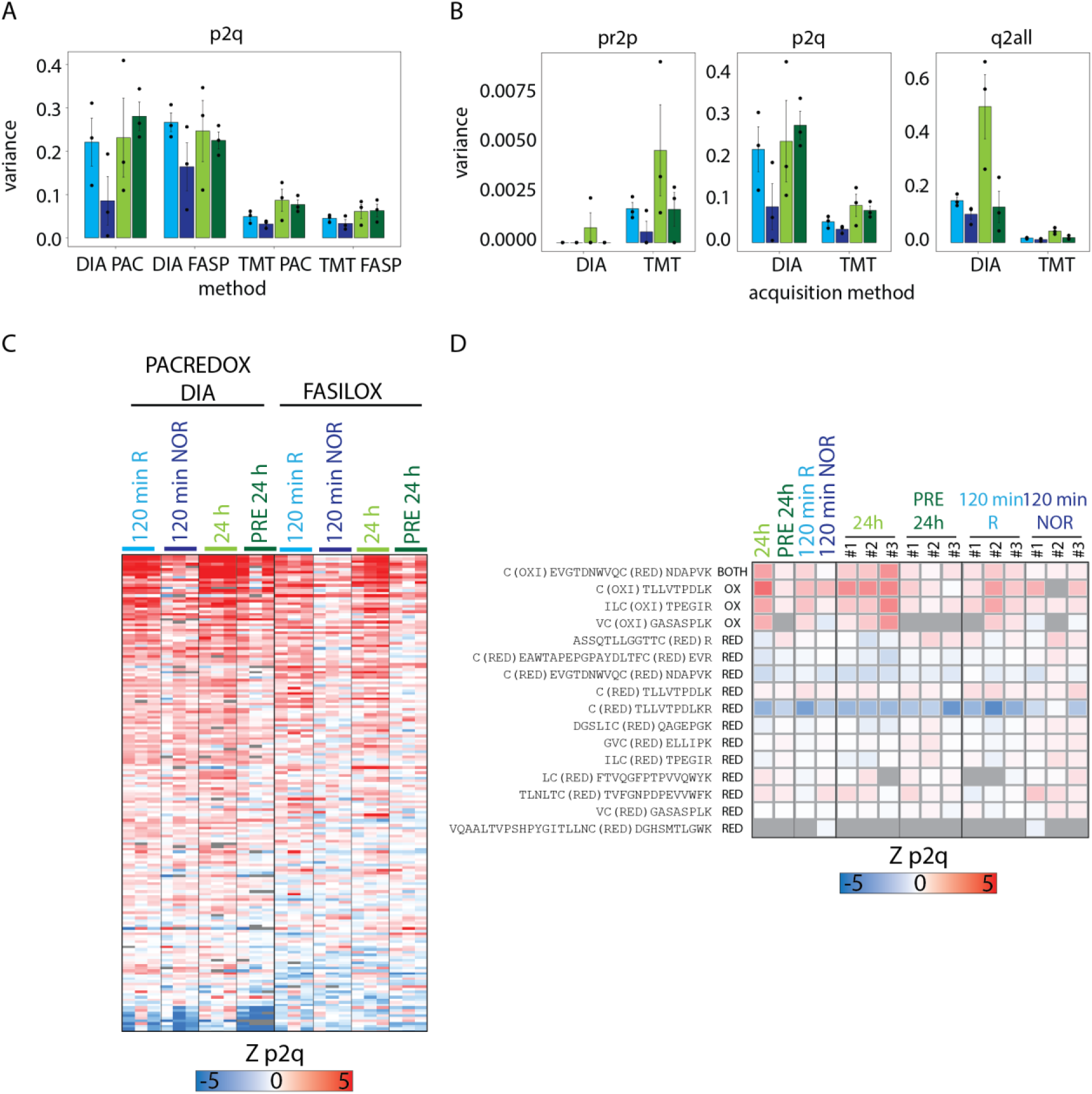
(A) Variances from four different experimental strategies (PAC-DIA, PAC-TMT, FASP-DIA, FASP-TMT) calculated in the peptide to protein integration level in iSanXot. Height of the bar is the average of n=3 biological replicates, and the error bar represents the standard deviation of the mean. Each colour indicates a different biological comparison (all of them are relative to baseline condition). (B) Variances from DIA or TMT quantification strategies calculated in the three integration levels (precursor or scan to peptide – pr2p, peptide to protein – p2q and protein to all – q2all) in iSanXot. Height of the bar is the average of n=3 biological replicates, and the error bar represents the standard deviation of the mean. Each colour indicates a different biological comparison (all of them are relative to baseline condition). (C) Heatmap of Zp2q values for MMTS-labelled peptides common in PACREDOX with DIA and FASILOX analysis. (D) Heatmap of Zp2q values for reduced and oxidized cysteine-containing peptides from MYOM2 protein detected exclusively by DIA analysis.

**Supplementary Figure 3.**
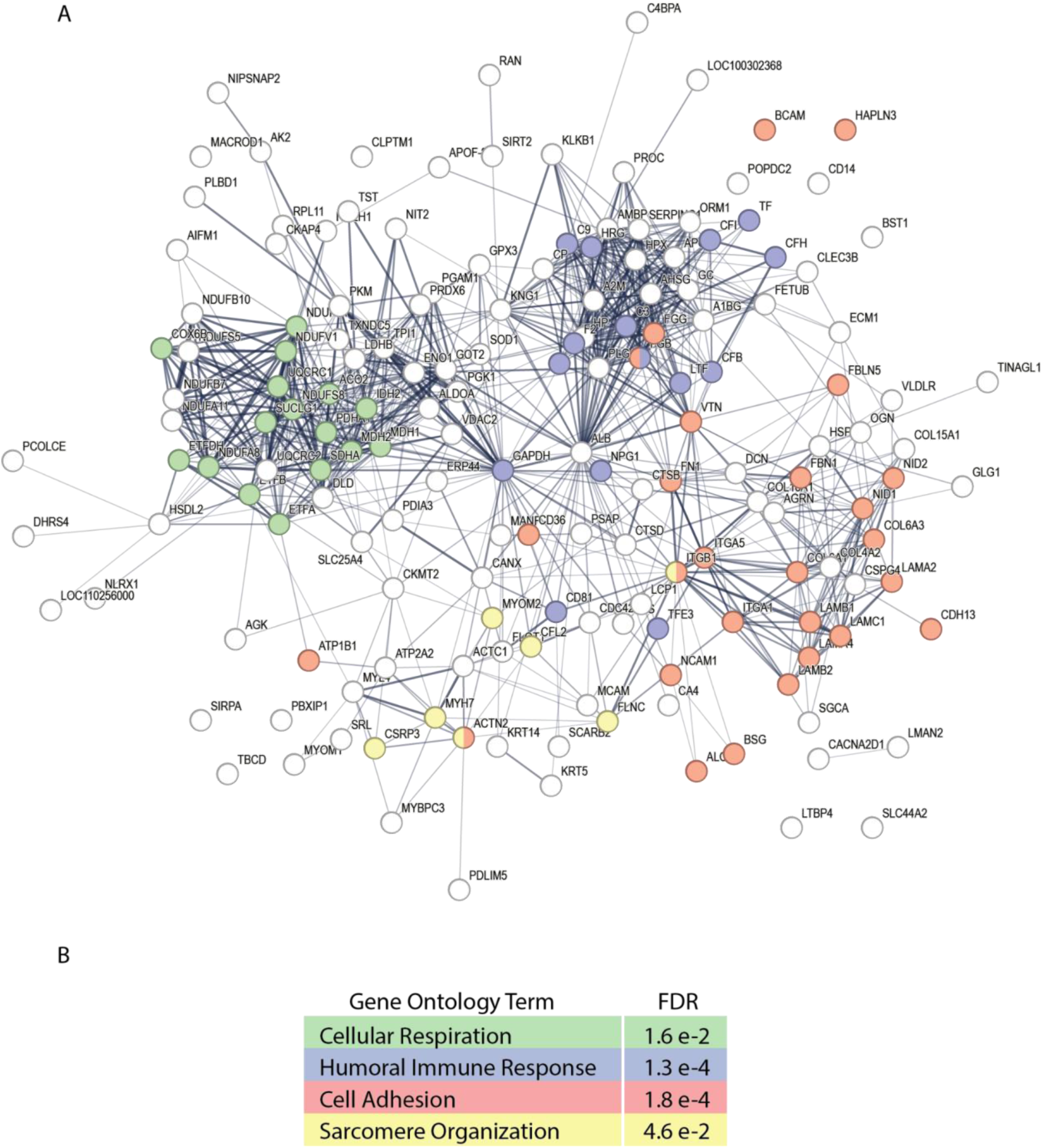
(A) STRING network of proteins containing reversibly oxidized cysteines detected in the PAC-redox DIA analysis. Colour represents whether the protein belongs to the Gene Ontology (GO) terms from panel B. (B) Most representative GO terms from the network shown in A, together with the FDR-corrected significance value for over-representation corrected by the measured proteomics background.

**Supplementary Figure 4.**
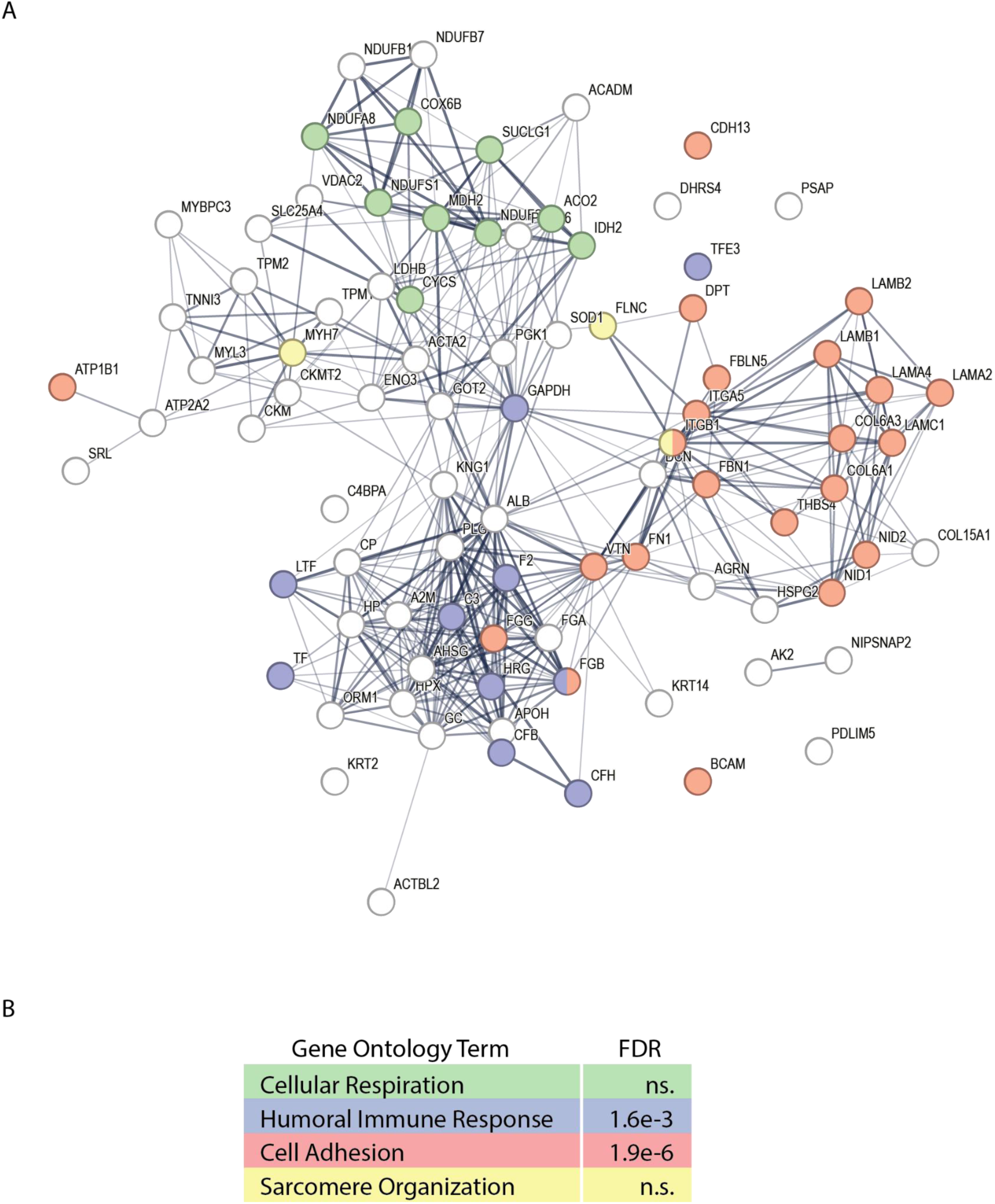
(A) STRING network of proteins containing reversibly oxidized cysteines detected in the FASILOX DIA analysis. Colour represents whether the protein belongs to the Gene Ontology (GO) terms from panel B. (B) Most representative GO terms from the network shown in A, together with the FDR-corrected significance value for over-representation corrected by the measured proteomics background. “n.s.” stands for not-significant.

**Supplementary Figure 5.**
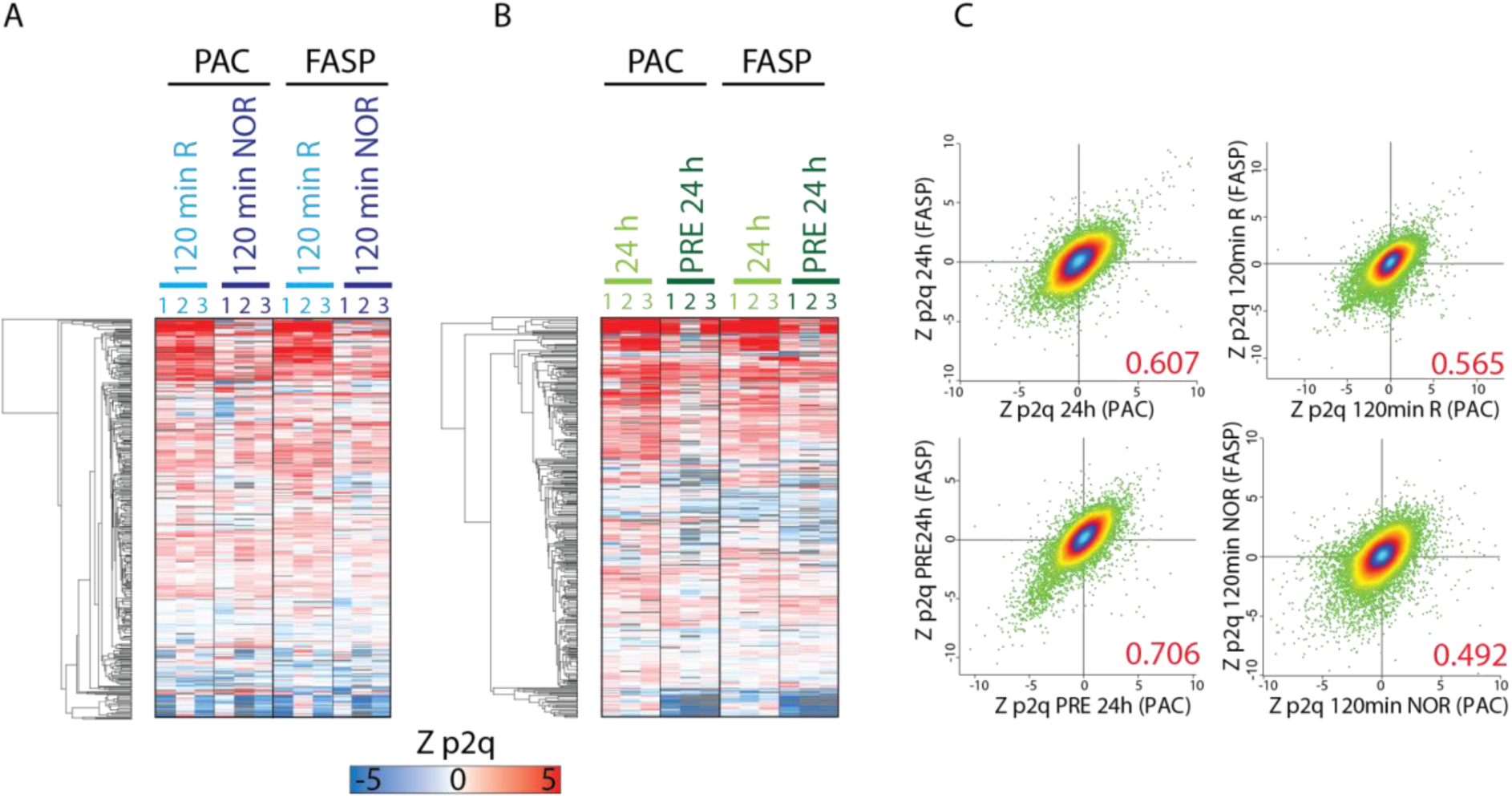
(A) Heatmap of Zp2q values for MMTS-labelled peptides common in FASP-redox and PAC-redox samples analysed using DIA for 120 min R and 120 min NOR samples (relative to baseline). (B) Heatmap of Zp2q values for MMTS-labelled peptides common in FASP-redox and PAC-redox samples analysed using DIA for 24 h and PRE 24 h samples (relative to baseline). (C) Correlation plots of Zp2q values calculated for all peptides in PAC versus FASP-based digestion experiments, both analysed using DIA. Colour indicates the density of the point distribution. Red value on the bottom right corner indicates the Person correlation of the distribution.

